# Distinct temporal requirements for Sonic hedgehog signaling in development of the tuberal hypothalamus

**DOI:** 10.1101/314773

**Authors:** Tanya S. Corman, Solsire E. Zevallos, Douglas J. Epstein

## Abstract

**ABSTRACT:** Sonic hedgehog (Shh) plays well characterized roles in the development of several regions of the brain and spinal cord, but its functions in the hypothalamus have been more difficult to elucidate due to the complex neuroanatomy of this brain area. Here, we utilize fate-mapping and conditional deletion models in mice to define requirements for dynamic Shh activity at distinct stages of tuberal hypothalamic development, a brain region with important homeostatic functions. At early time points, Shh signaling regulates dorsoventral patterning, neurogenesis, and the size of the ventral midline. Fate mapping experiments demonstrate that Shh expressing and responsive progenitors contribute to distinct neuronal subtypes, accounting for some of the cellular heterogeneity in tuberal hypothalamic nuclei. Conditional deletion of the Hedgehog transducer Smoothened (Smo), after dorsoventral patterning has been established, reveals that Shh signaling is necessary to maintain proliferation and progenitor identity during peak periods of hypothalamic neurogenesis. We also find that mosaic disruption of *Smo* causes a non-cell autonomous gain in Shh signaling activity in neighboring wild type cells, suggesting a mechanism for the pathogenesis of hypothalamic hamartomas, a benign tumor that forms during hypothalamic development.

**SUMMARY STATEMENT:** Requirements for dynamic Sonic hedgehog activity at distinct stages of tuberal hypothalamic development are defined using fate-mapping and conditional deletion models in mice.

## INTRODUCTION

The hypothalamus is an ancient brain region with important roles in the regulation of several homeostatic processes and animal behaviors. Neural circuits mapping to distinct areas of the hypothalamus control a variety of essential bodily functions, including food intake, energy expenditure, fluid balance, temperature regulation, wakefulness, daily rhythms, as well as social behaviors associated with reproduction, aggression, arousal, and stress (Saper and Lowell, 2014; Zha and Xu, 2015; Hashikawa et al., 2017; Tan and Knight, 2018). Organized into small clusters of neurons, termed nuclei, the hypothalamus is unlike other regions of the central nervous system (CNS) that are typically arranged in cell layers (Shimada and Nakamura, 1973; Altman and Bayer, 1986). Further adding to this complex architecture, most hypothalamic nuclei are composed of diverse neuronal cell types with opposing or sometimes unrelated functions. The developmental mechanisms regulating neuronal heterogeneity within hypothalamic nuclei are poorly understood compared to other CNS regions (Bedont et al., 2015; Burbridge et al., 2016; Xie and Dorsky, 2017).

The hypothalamus derives from the ventral diencephalon and can be divided into four principal regions from rostral to caudal: preoptic, anterior, tuberal, and mammillary. Within the tuberal hypothalamus, neurons in the arcuate nucleus (ARC), ventromedial hypothalamic nucleus (VMH), dorsomedial hypothalamic nucleus (DMH), and paraventricular nucleus (PVN) integrate sensory information from the environment in order to illicit autonomic, endocrine and behavioral responses that maintain vital set points in the animal or adapt to various stressors (Saper and Lowell, 2014). In addition to receiving inputs from other brain regions, the positioning of the tuberal hypothalamus offers unique exposure to peripheral cues carried through the bloodstream which are presented to neurons at the median eminence.

Recent advances have been made in our understanding of the neuronal circuitry and function of tuberal hypothalamic nuclei. This is particularly true for the VMH, an elliptical shaped nucleus located above the ARC and below the DMH (Fig. 1). The VMH is subdivided into ventrolateral (VMH_VL_), central (VMH_C_) and dorsomedial (VMH_DM_) regions, each with distinguishable gene expression profiles (Segal et al., 2005; McClellan et al., 2006; Kurrasch et al., 2007). Genetic, pharmacogenetic and optogenetic approaches have further delineated VMH neurons into functionally distinct categories. ERα-expressing neurons in the VMH_VL_ regulate sexually dimorphic features related to energy expenditure, reproductive behavior and aggression (Lin et al., 2011; Lee et al., 2014; Correa et al., 2015). By contrast, insulin receptor (IR) expressing neurons in the VMH_C_ and Leptin Receptor (LEPR) neurons in the VMH_DM_ effect body weight regulation in both males and females (Dhillon et al., 2006; Klöckener et al., 2011). Steroidogenic factor 1 (SF1, officially designated Nr5a1) neurons in the VMH_DM_ also influence behavioral responses to fear and anxiety (Silva et al., 2013; Kunwar et al., 2015).

**Figure 1.**
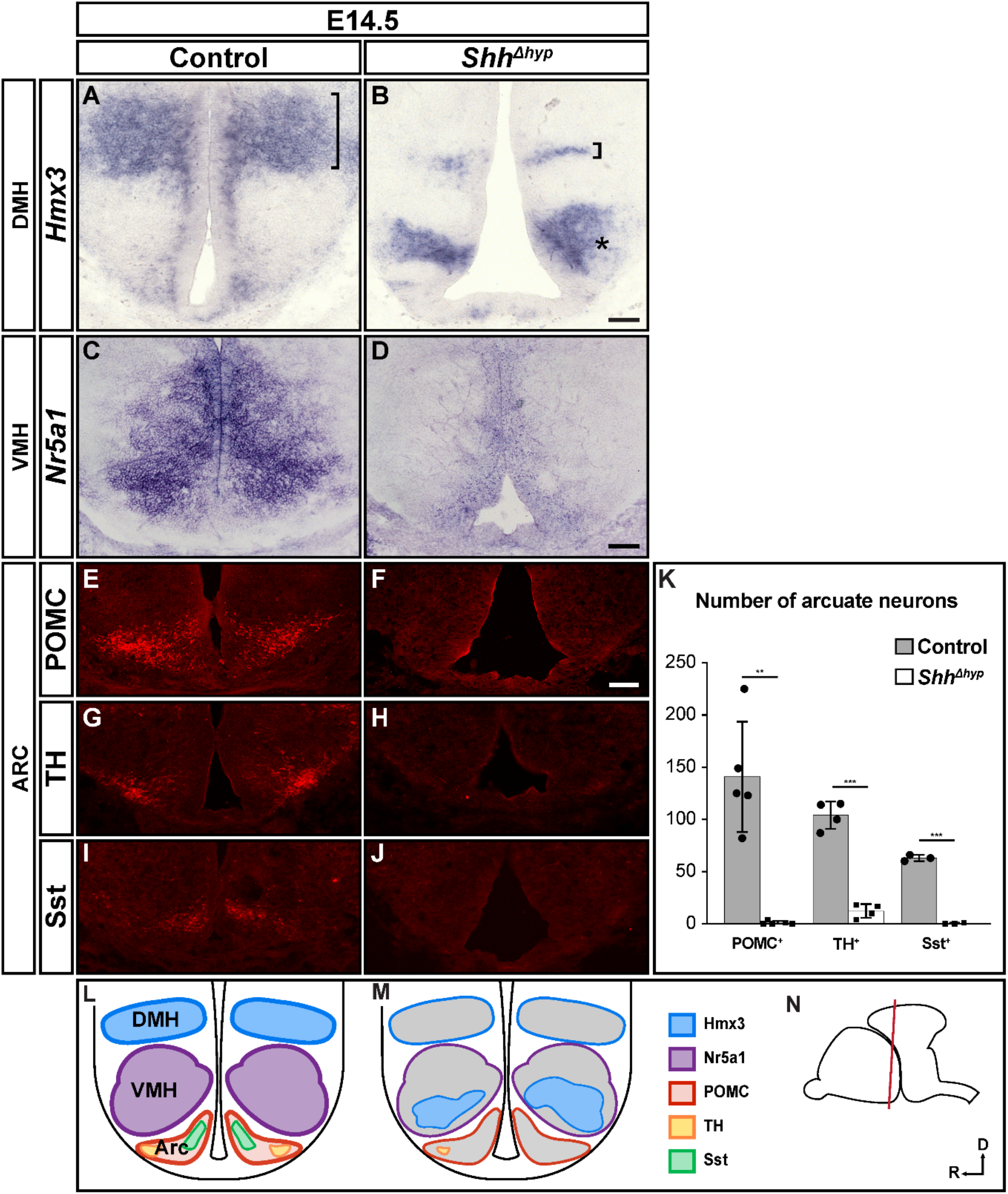
Loss of tuberal hypothalamic neurons in *Shh^Δhyp^* embryos. Coronal sections through the tuberal hypothalamus of control and *Shh^Δhyp^* embryos stained by RNA *in situ* hybridization (A-D) or immunofluorescence (E-J) at E14.5 for neuronal markers. (A,B) *Hmx3* is expressed in the DMH of control embryos, and shows reduced (bracket) and ectopic (asterisk) expression in *Shh^Δhyp^* mutants (n=3). (C,D) *Nr5a1* marks the VMH in control embryos and is absent in *Shh^Δhyp^* mutants (n=4). (E-K) The number of cells expressing markers of distinct neuronal subtypes in the ARC (POMC, TH, and Sst) are reduced in *Shh^Δhyp^* mutants (n=5 for POMC, ^**^p=0.0004; n=4 for TH, ^***^p<0.0001; n= 3 for Sst, ^***^p<0.0001). (L,M) Schematic of coronal sections through the tuberal hypothalamus of control and *Shh^Δhyp^* embryos showing nuclei and cell type specific markers. (N) Sagittal view of brain showing plane of section (red line) in (L-M). Scale bars: 100 μm. Error bars indicate S.D. Statistical analysis was performed using a two-tailed unpaired *t*-test.

Despite the progress in assigning functions to VMH neurons, we still know relatively little about how this nucleus forms and the process by which its subdivisions are established. During hypothalamic development, Nr5a1 is selectively expressed by all VMH neurons soon after they exit the cell cycle and undergo neurogenesis (Tran et al., 2003). Nr5a1 is required for the terminal differentiation of VMH neurons, as well as their coalescence into a nucleus with a distinct cytoarchitecture (Ikeda et al., 1995; Davis et al., 2004; Büdefeld et al., 2011). Consequently, mice lacking Nr5a1 in the VMH are obese, anxious, and infertile (Majdic et al., 2002; Zhao et al., 2008; Kim et al., 2010). Additional cell type specific factors acting upstream of Nr5a1 remain to be identified.

One signaling molecule that may help bridge the gap in knowledge concerning the ontogeny of VMH neurons is Sonic Hedgehog (Shh). Shh has been studied in several spatial and temporal contexts related to hypothalamic development. Shh signaling from the prechordal plate, which underlies the ventral forebrain at early stages of its development, is required for the induction of the hypothalamic territory (Chiang et al., 1996; Dale et al., 1997). Conditional deletion of Shh in the ventral diencephalon causes defects in the patterning, regionalization and formation of ventral hypothalamic nuclei (Szabó et al., 2009; Shimogori et al., 2010; Zhao et al., 2012; Carreno et al., 2017). Nevertheless, the pathogenic mechanisms underlying these Shh dependent alterations in hypothalamic development have yet to be fully elucidated. Moreover, since Shh continues to be expressed in VMH progenitors well beyond the initial patterning stage, additional roles for Shh in VMH nucleogenesis and neuronal subtype identity are likely (Alvarez-Bolado et al., 2012).

Here, we use conditional knockout mice to interrogate the functional requirements for Shh signaling at specific periods of hypothalamic development. We show that the pronounced loss of hypothalamic nuclei that manifests from early deletion of Shh at embryonic day 9 (E9.0) is caused by defects in dorsoventral patterning, neurogenesis and the expansion of ventral midline cells, indicating a novel role for Shh in restricting ventral midline development in the tuberal hypothalamus. Fate mapping experiments reveal that Shh expressing and Shh responsive cell lineages are enriched in distinct domains of the VMH, contributing to the neuronal heterogeneity of this nucleus. Deletion of Smoothened (Smo), an essential transducer of Shh signaling, at later stages of hypothalamic development (after E10.5), resulted in a cell autonomous loss of VMH neuronal subtype identity. Remarkably, we also detect a non-cell autonomous expansion and reprogramming of neighboring wild type cells, which likely occurred in response to residual Shh ligand that was not taken up by *Smo* mutant cells. This gain in Shh signaling activity may explain the pathogenesis of hypothalamic hamartomas (HH), a benign tumor caused, in some cases, by somatic gene mutations that block Shh responsiveness (Saitsu et al. 2016; Hildebrand et al., 2016).

## RESULTS

### Shh is required for development of tuberal hypothalamic nuclei

To determine how Shh signaling contributes to formation of tuberal hypothalamic nuclei we first evaluated the expression of cell type specific markers in *SBE2-cre; Shh^loxp/-^* (*Shh^Δhyp^*) embryos. *SBE2-cre* is a transgenic mouse line that uses Shh brain enhancer 2 (SBE2) to activate *cre* transcription in the ventral diencephalon in a similar pattern to the endogenous expression of *Shh*. We previously showed that Shh is selectively deleted in the ventral diencephalon of *Shh^Δhyp^* embryos by E9.0 (Zhao et al., 2012). Expression of cell type specific markers of the DMH (Hmx3), VMH (Nr5a1) and ARC (Pro-opiomelanocortin, POMC; Tyrosine Hydoxylase, TH; and Somatostatin, Sst) nuclei was either absent or greatly diminished in *Shh^Δhyp^* embryos at E14.5 (Fig. 1A-K; POMC: control 140.8 ± 52.9, *Shh^Δhyp^* 1.0 ± 1.7, n=5, p=0.0004; TH: control 104.0 ± 13.2, *Shh^Δhyp^* 12.3 ± 6.6, n=4, p<0.0001; Sst: control 63.0 ± 3.0, *Shh^Δhyp^* 0.3 ± 0.6, n=3, p<0.0001). Ectopic expression of *Hmx3* was also detected in the VMH, possibly due to its derepression in the absence of Shh (Fig. 1A, B). These results are consistent with previous findings demonstrating a requirement for Shh in the development of tuberal hypothalamic nuclei (Fig. 1L-N; Szabó et al., 2009; Shimogori et al., 2010; Carreno et al., 2017).

### Alterations in dorsoventral patterning, neurogenesis and ventral midline formation explain the absence of tuberal hypothalamic nuclei in *Shh^Δhyp^* embryos

Shh signaling is required to establish distinct neuronal identities at ventral positions along the length of the vertebrate neural tube through activation and repression of homeodomain and basic helix-loop-helix (bHLH) transcription factors (Ericson et al., 1997; Briscoe et al., 1999; Briscoe et al., 2000; Muhr et al., 2001; Balaskas et al., 2012). However, the temporal and spatial dynamics of Shh signaling in the hypothalamus differ from those in more posterior regions of the CNS. *Shh* is initially broadly expressed in ventral hypothalamic progenitors and then rapidly downregulated in the ventral midline at the level of the tuberal hypothalamus (Manning et al., 2006; Trowe et al., 2013). As a result, *Shh* is expressed in bilateral stripes adjacent to the ventral midline, unlike in spinal cord and hindbrain regions where *Shh* is restricted to the floor plate (Fig. 2A; and Echelard et al., 1993). Moreover, neural progenitors immediately dorsal to the bilateral stripes of *Shh* are responsive to Shh signaling, indicated by *Gli1* expression, whereas progenitors located in the ventral midline are refractory to Shh signaling (Fig. 2B; and Ohyama et al., 2008).

**Figure 2.**
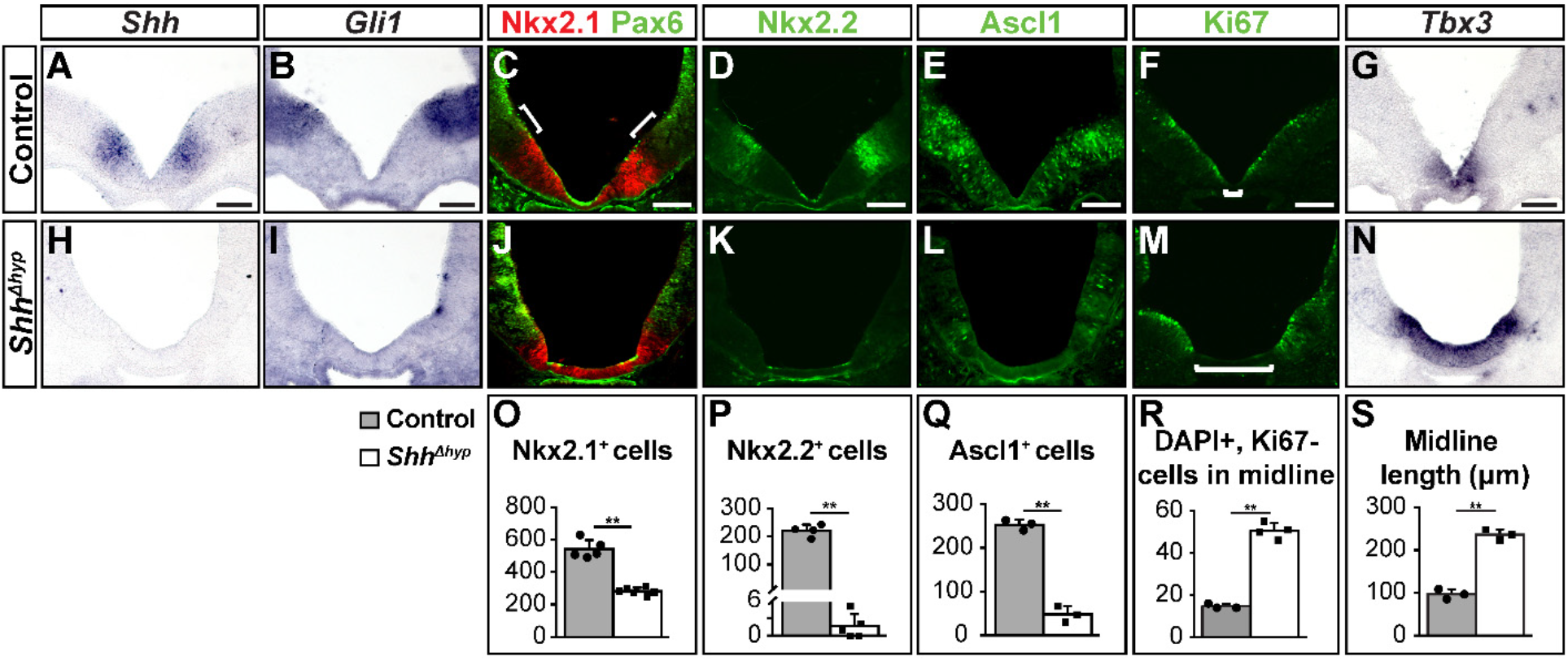
Altered dorsoventral patterning, neurogenesis, and ventral midline development in *Shh^Δhyp^* embryos. Coronal sections through control and *Shh^Δhyp^* embryos stained by RNA *in situ* hybridization (A,B,G-I,N) or immunofluorescence (C-F,J-M) at E10.5. (A,H) *Shh* expression in the prospective tuberal hypothalamus is lost in *Shh^Δhyp^* mutants (n=3). (B,I) *Gli1* expression in neuronal progenitors immediately dorsal to *Shh* is absent in *Shh^Δhyp^* mutants (n=3). (C,J,O) The number of cells in the ventral hypothalamus expressing Nkx2.1 is reduced in *Shh^Δhyp^* mutants (n=5 for controls, n=6 for *Shh^Δhyp^*), whereas the domain of Pax6 expression is expanded ventrally. Note that the gap between Nkx2.1 and Pax6 (white bracket) in control embryos is missing in *Shh^Δhyp^* mutants. (D,K,P) The number of Nkx2.2 expressing cells in tuberal hypothalamic progenitors is greatly reduced in *Shh^Δhyp^* mutants (n=4 for controls, n=5 for *Shh^Δhyp^*). (E,L,Q) The number of Ascl1 expressing neurogenic progenitors is reduced in *Shh^Δhyp^* mutants (n=3). (F,M,R) The number of non-proliferating ventral midline cells identified by the absence of Ki67 staining is expanded in *Shh^Δhyp^* mutants (n=4). (G,N,S) The expression of *Tbx3* in the ventral midline is expanded in *Shh^Δhyp^* mutants (n=3). Scale bars: 100 μm. Error bars indicate S.D. Statistical analysis was performed using a two-tailed unpaired *t*-test (^**^p<0.0001).

We evaluated key patterning genes in *Shh^Δhyp^* mutants at E10.5 to address whether alterations in their expression might explain the loss of tuberal hypothalamic nuclei at later stages. As expected, *Shh* and *Gli1* were absent in *Shh^Δhyp^* embryos (Fig. 2A,B,H,I). The number of cells expressing Nkx2.1, a broad determinant of ventral hypothalamic progenitor identity (Kimura et al., 1996; Sussel et al., 1999), was reduced by 48% in *Shh^Δhyp^* embryos compared to control littermates (Fig. 2C,J,O; control: 540.8 ± 56.4, n=5; *Shh^Δhyp^*: 282.0 ± 22.0, n=6, p<0.0001). In addition, Pax6, a prethalamic marker was expanded ventrally to a position abutting the Nkx2.1 domain (Fig. 2C,J). Nkx2.2 is expressed in Shh responsive progenitors of the DMH that occupy the gap between Pax6 – and Nkx2.1 – positive cells (Fig. 2D). Expression of Nkx2.2 was dramatically reduced in *Shh^Δhyp^* embryos (Fig. 2D,K,P; control: 221.0 ± 21.2, n=4; *Shh^Δhyp^*: 1.6 ± 2.1, n=5, p<0.0001), which may explain the ventral expansion of Pax6 due to deficient cross-repressive interactions between these transcription factors (Ericson et al., 1997; Briscoe et al., 1999; Muhr et al., 2001). These results suggest that alterations in the dorsoventral patterning of tuberal hypothalamic progenitors are responsible, at least in part, for the loss of their identities in *Shh^Δhyp^* embryos.

The bHLH transcription factor, Ascl1, is required for ARC and VMH neurogenesis (McNay et al., 2006). We observed an 81% reduction in the number of Ascl1 expressing cells in the ventral hypothalamus of *Shh^Δhyp^* embryos compared to control littermates (Fig. 2E,L,Q; control: 252.7 ± 11.7, n=3; *Shh^Δhyp^*: 47.0 ± 18.5, n=3, p<0.0001). Therefore, the reduction of neuroendocrine neurons in the ARC and VMH nuclei of *Shh^Δhyp^* embryos may also be explained by the downregulation of Ascl1.

Another striking feature of *Shh^Δhyp^* embryos is the dysmorphic appearance of the ventral midline. Rather than the V-shaped morphology typical of wild type embryos, the ventral midline of *Shh^Δhyp^* embryos is U-shaped with a flattened appearance (Fig. 2F,M). Ventral midline cells in the tuberal hypothalamus undergo BMP dependent cell-cycle arrest and express the T-box protein, Tbx3 (Manning et al., 2006; Trowe et al., 2013). We quantified the zone of non-proliferation by counting the number of DAPI positive nuclei in the Ki67 negative ventral midline territory and observed a significant increase in *Shh^Δhyp^* embryos (Fig. 2F,M,R; 50.5 ± 3.7, n=4) compared to control littermates (14.67 ± 1.5, n=3, p<0.0001). The length of the ventral midline territory marked by Tbx3 was also significantly increased in *Shh^Δhyp^* embryos (Fig. 2G,N,S; 239.1 ± 8.2 μm, n=3) compared to control littermates (96.7 ± 11.6 μm, n=3, p=0.0001). These results demonstrate that the loss of Shh in the tuberal hypothalamus expands the fate of non-dividing ventral midline cells at the expense of proliferating Nkx2.1 positive neural progenitors. Interestingly, *Bmp4* was previously shown to be upregulated in the ventral midline of *Shh^Δhyp^* embryos (Zhao et al., 2012). Hence, our findings highlight a unique role for Shh in restricting the size of the ventral midline in the tuberal hypothalamus by opposing Bmp signaling, in stark contrast to the floor plate promoting activity of Shh in more posterior regions of the CNS (Echelard et al., 1993; Roelink et al., 1994; Roelink et al., 1995; Marti et al., 1995; Ericson et al., 1996; Chiang et al., 1996).

### Descendants of *Shh* expressing progenitors contribute to the VMH

*Shh* and *Gli1* continue to be expressed beyond the stage when tuberal hypothalamic progenitors first acquire their identity and extend into the peak period of DMH, VMH and ARC neurogenesis between E12.5 and E14.5 (Fig. S1). We next sought to determine the relative contributions of Shh expressing and Shh responsive progenitors to distinct tuberal hypothalamic nuclei. We used a genetic fate mapping strategy to indelibly label Shh expressing (*Shh^CreER^*) and Shh responsive (*Gli1^CreER^*) progenitors with an inducible GFP reporter (*Rosa^ZsGreen^*). Pregnant dams carrying either *Shh^CreER/+^; Rosa^ZsGreen/+^* or *Gli1^CreER/+^; Rosa^ZsGreen/+^* embryos were administered tamoxifen at E10.5. The contribution of *Shh* and *Gli1* expressing cells to tuberal hypothalamic nuclei was evaluated by co-labeling with GFP and cell type specific markers at E14.5.

The majority of postmitotic neurons in the VMH express Nr5a1 and Nkx2.1 at E14.5 (Fig. 3A,B; Correa et al., 2015). GFP^+^ cells in *Shh^CreER/+^; Rosa^ZsGreen/+^* embryos were detected in the ventricular zone adjacent to the VMH as well as 46% of Nr5a1 positive neurons (Fig. 3A,G). This finding suggests that Shh expressing progenitors in the ventricular zone of the tuberal hypothalamus migrate radially to populate the VMH, in agreement with a previous study (Alvarez-Bolado et al., 2012). GFP and Nr5a1 double labeled neurons were enriched in ventral (199.3 ± 11.9, n=3) compared to dorsal (112.3 ± 8.0, n=3, p=0.0005) regions of the VMH in *Shh^CreER/+^; Rosa^ZsGreen/+^* embryos (Figs. 3A and S2). A similar ventral VMH bias was observed for GFP and Nkx2.1 double labeled neurons (Fig. 3B). Nkx2.2 is expressed in a subset of post-mitotic neurons in the dorsal VMH, of which 22% co-label with GFP (Fig. 3C,G). These data reveal that descendants of Shh expressing progenitors marked with GFP at E10.5 are partitioned along the dorsoventral axis of the VMH into primarily ventral and central positions and are largely excluded from the dorsal most region (Figs. 3J and S2).

**Figure 3.**
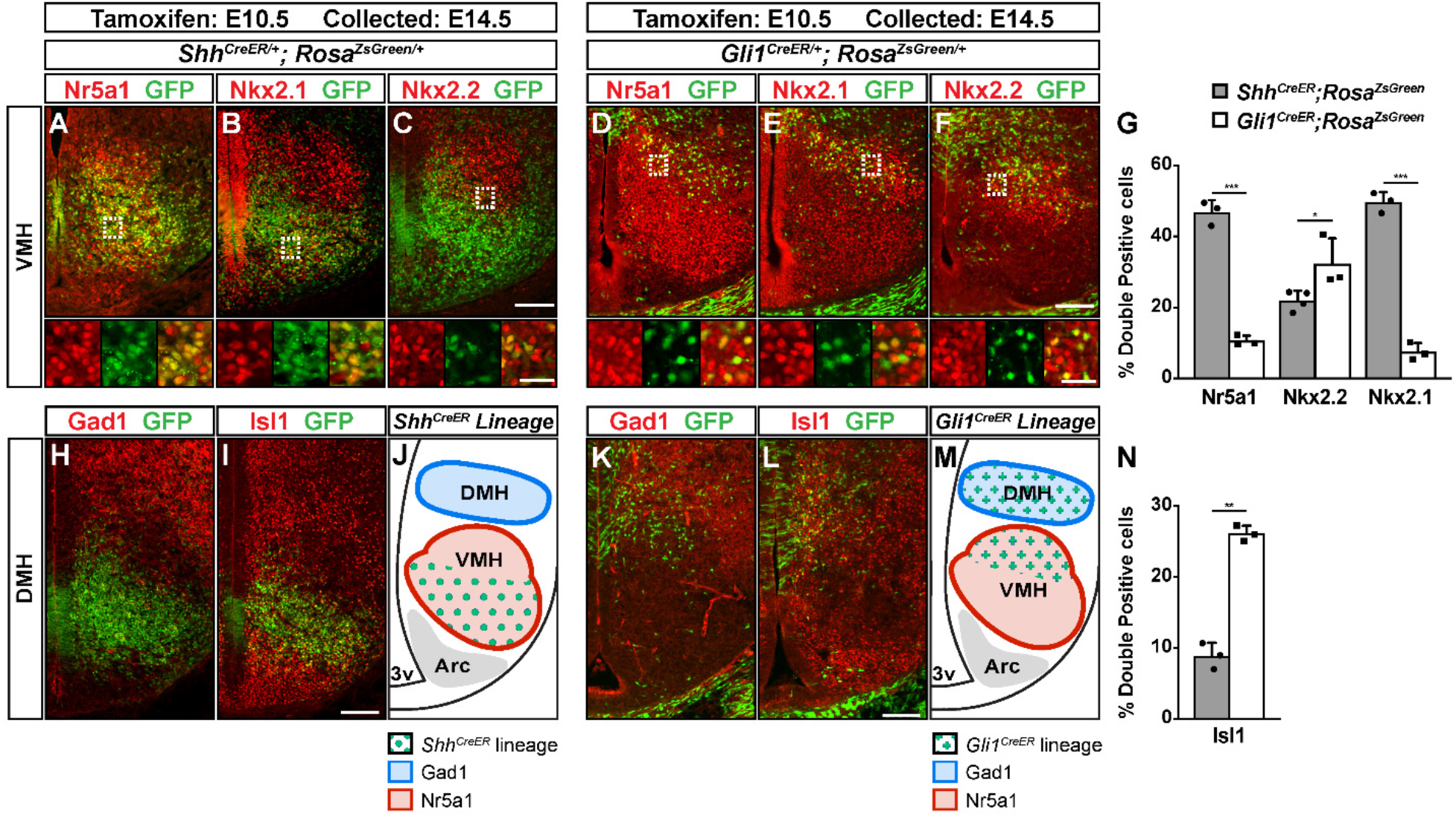
*Shh* and *Gli1* expressing cells contribute to overlapping and distinct tuberal hypothalamic nuclei. Coronal sections through the tuberal hypothalamus of *Shh^CreER/+^;Rosa^ZsGreen/+^* (A-C,H, I) and *Gli1^CreER/+^; Rosa^ZsGreen/+^* (D-F,K,L) embryos at E14.5 that received tamoxifen at E10.5. The fate of *Shh* and *Gli1* expressing cells was determined by co-labeling with GFP (ZsGreen) and cell type specific markers of the VMH and DMH. Insets in (A-F) are higher magnification views of boxed regions showing single and double labeled cells. (G,N) Quantification of fate-mapping experiments displayed as the proportion of double positive cells (from either Shh or Gli1 lineages) to the total number of cells expressing a given marker: Nr5a1(^***^p<0.0001, n=3); Nkx2.2 (^*^p=0.038, n=4 for Shh lineage, n=3 for Gli1 lineage); Nkx2.1 (^***^p<0.0001, n=3); Isl1 (^**^p<0.0003, n=3). (J,M) Schematic demonstrating contribution of *Shh* and *Gli1* expressing cell lineages to tuberal hypopthalamic nuclei. Scale bars: 100 μm. Inset scale bars: 25 μm. Error bars indicate S.D. Statistical analysis was performed using a two-tailed unpaired *t*-test on arcsin-transformed data.

We also assessed the contribution of Shh expressing progenitors to other tuberal hypothalamic nuclei. The DMH is composed of GABA-ergic neurons that express Gad1 and a subset of neurons that express Isl1. No overlap in GFP and Gad1 or Isl1 was observed in the DMH of *Shh^CreER/+^; Rosa^ZsGreen/+^* embryos, suggesting that Shh expressing progenitors do not contribute to this nucleus when labeled at E10.5 (Fig. 3H-J). Some overlap between GFP and Isl1 was observed in ventrolateral neurons in the VMH, as well as a small subset of ARC neurons (Fig. 3I,N). In summary, Shh expressing neuronal progenitors in the tuberal hypothalamus predominantly contribute to the VMH when labeled at E10.5.

### Descendants of *Gli1* expressing progenitors contribute to the DMH and VMH

We next examined the fate of Shh responsive cells in *Gli1^CreER/+^; Rosa^ZsGreen/+^*embryos administered tamoxifen at E10.5 and collected at E14.5. Given that *Gli1* is expressed dorsal to *Shh*, we expected that GFP^+^ cells would be confined to the DMH. Indeed, GFP^+^ cells were detected in the ventricular zone immediately adjacent to the DMH, as well as in Gad1 and Isl1 expressing neurons that appeared to migrate radially into the DMH (Fig. 3K-N).

Surprisingly, we also identified a small population of Nr5a1 neurons in the VMH of *Gli1^CreER/+^; Rosa^ZsGreen/+^* embryos that co-labeled with GFP (Fig. 3D,G; 10.6% ± 1.4%, n=3). These GFP^+^ cells appeared to stream ventrally from a more dorsal progenitor domain to occupy a dorsolateral region of the VMH (Fig. 3D-F). GFP and Nr5a1 double labeled neurons were enriched in dorsal (57.3 ± 13.5, n=3) compared to ventral (5.0 ± 1.7, n=3, p=0.002) regions of the VMH in *Gli1^CreER/+^; Rosa^ZsGreen/+^* embryos (Figs. 3D and S2). GFP co-labeling was also observed with Nkx2.1 and Nkx2.2 in primarily dorsal regions of the VMH (Fig. 3E-G). These data suggest that VMH neurons originate from spatially segregated pools of Shh expressing and Shh responsive progenitors that may contribute to the cellular heterogeneity of the VMH (Fig.3J,M).

### Shh signaling is required to maintain tuberal hypothalamic progenitors in a proliferative state

The results of our lineage tracing and gene expression experiments demonstrated that Shh signaling is active in tuberal hypothalamic progenitors after their dorsoventral identity is established. To address whether Shh signaling is also required at later stages of DMH and VMH neurogenesis we generated mice in which *Smoothened* (*Smo*), an essential regulator of Shh signaling, was conditionally deleted in *Gli1* expressing cells after E10.5.

Pregnant dams carrying *Gli1^CreER/+^*; *Smo^loxp/-^; Rosa^ZsGreen/+^* (*cSmo*) and *Gli1^CreER/+^*; *Smo^loxp/+^; Rosa^ZsGreen/+^* (control) embryos were administered tamoxifen at E10.5 and collected at different developmental stages. Strikingly, the number of GFP positive cells in the tuberal hypothalamus was decreased by 43% in *cSmo* embryos (453.8 ± 52.27, n=8) compared to control littermates (800.2 ± 155.3, n=9, p=0.0003) when harvested at E14.5 (Fig. 4A-C). This reduction in GFP^+^ cells was detected in both mantle and ventricular zones. A significant reduction in the number of GFP^+^ cells was also observed one day earlier at E13.5 (Fig. 4D-F; *cSmo*: 418.1 ± 45.5, n=7, versus control: 603 ± 138.9, n=6, p=0.0066), but not at E12.5 (Fig. G-I; *cSmo*: 512.4 ± 96.1, n=5, versus control: 578.6 ± 96.7, n=5, p>0.05).

The failed expansion of Shh responsive cells in *cSmo* embryos between E12.5 and E14.5 might be explained by increased cell death, decreased proliferation, or both. Immunostaining for activated caspase-3, an indicator of apoptosis, was minimal in *cSmo* and control embryos at E14.5 with no significant differences between genotypes (Fig. S3). On the other hand, fewer GFP positive cells co-labeled with the proliferation marker Ki67 in the ventricular zone of *cSmo* embryos at E12.5 (Fig. 4J-L; 28.5% ± 6.2%, n=3) compared to control littermates (50.1% ± 8.7%, n=9, p=0.0249). From these results, we conclude that Shh signaling is required to maintain tuberal hypothalamic progenitors in a mitotically active state during the peak period of VMH and DMH neurogenesis.

**Figure 4.**
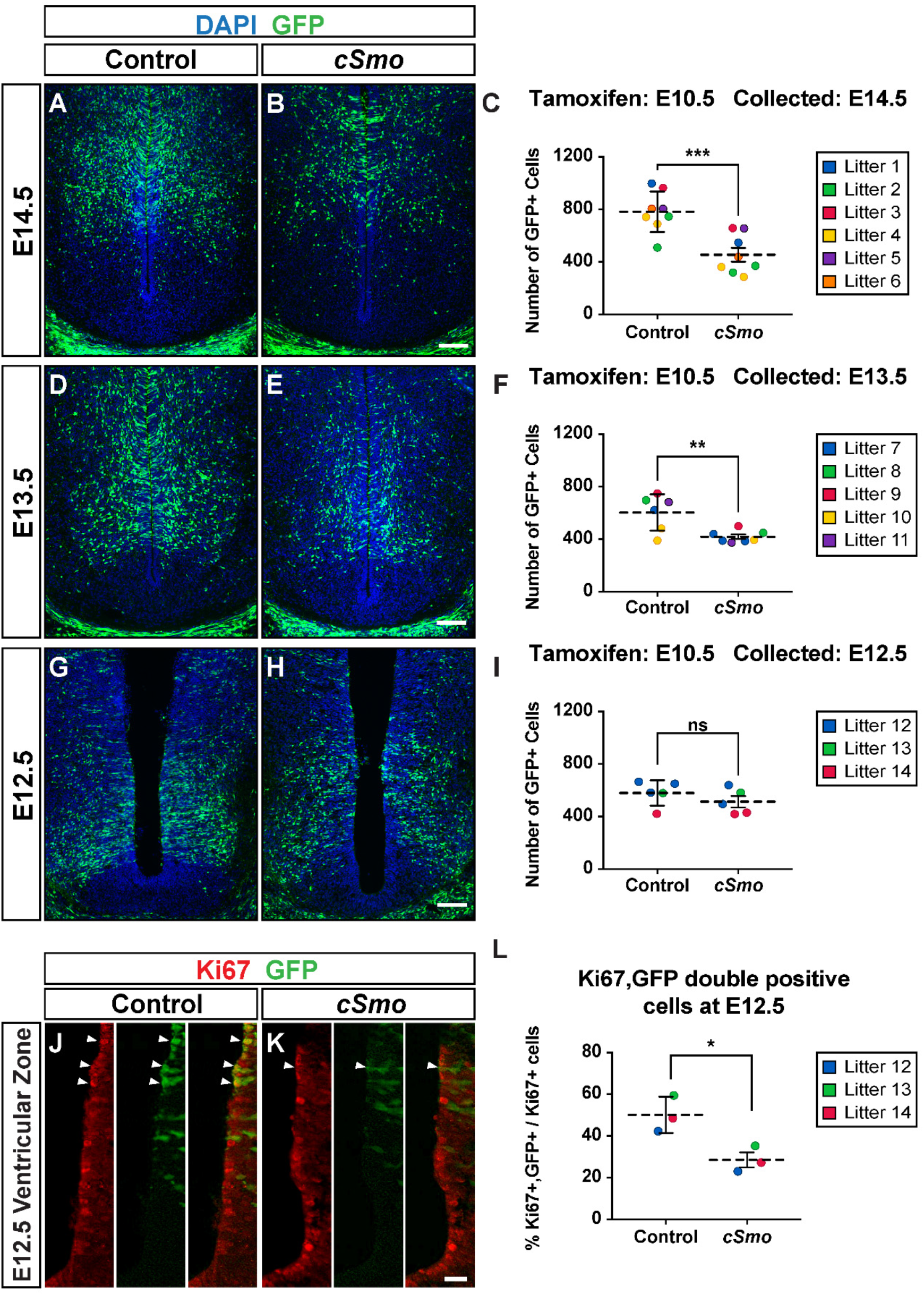
Shh signaling is required for proliferation of tuberal hypothalamic progenitors. Coronal sections through the tuberal hypothalamus of control and *cSmo* embryos at E14.5 (A,B), E13.5 (D,E), and E12.5 (G,H,J,K) that received tamoxifen at E10.5. (A-H) GFP (ZsGreen) fluorescence is shown on sections counterstained with DAPI. (C,F,I) Quantification of the number of GFP positive cells from *cSmo* and control embryos tracked by litter. (J-K) Co-labeling of GFP and Ki67 (arrowhead) in the ventricular zone reveals a reduction in the number of proliferating GFP positive cells in *cSmo* embryos at E12.5, as quantified in (L). Scale bars: (A-H) 100 μm (J-K) 25 μm. For all graphs, horizontal dotted line represents the mean and error bars indicate S.D. Each data point represents a single embryo that is color-coded by litter. Statistical analysis was performed using a two-tailed unpaired *t*-test (^***^p=0.0003, ^**^p=0.0066, ^*^p=0.0249).

### Cell-autonomous requirement for Smo in promoting tuberal hypothalamic neuron identity

Given the contribution of Shh responsive cells to distinct neuronal subtypes in the DMH and VMH, we next assessed whether aspects of their identity were compromised in *cSmo* mutants. We observed a 30% reduction in the number of Isl1 positive neurons in the DMH of *cSmo* (Fig. 5A-C; 477.6 ± 130.7, n=5) compared to control littermates (678.3 ± 44.6, n=4, p=0.0229). There was also a 30% reduction in the number of neurons expressing Nkx2.2 in the dorsal VMH of *cSmo* (Fig. 5D-F; 315.2 ± 52.1, n=9) compared to control embryos (451.6 ± 28.2, n=9; p<0.0001). These findings, in conjunction with the proliferation defects described above, suggest that the conditional loss of Shh signaling after E10.5 causes a cell autonomous reduction of distinct DMH and VMH neuronal subtypes.

**Figure 5.**
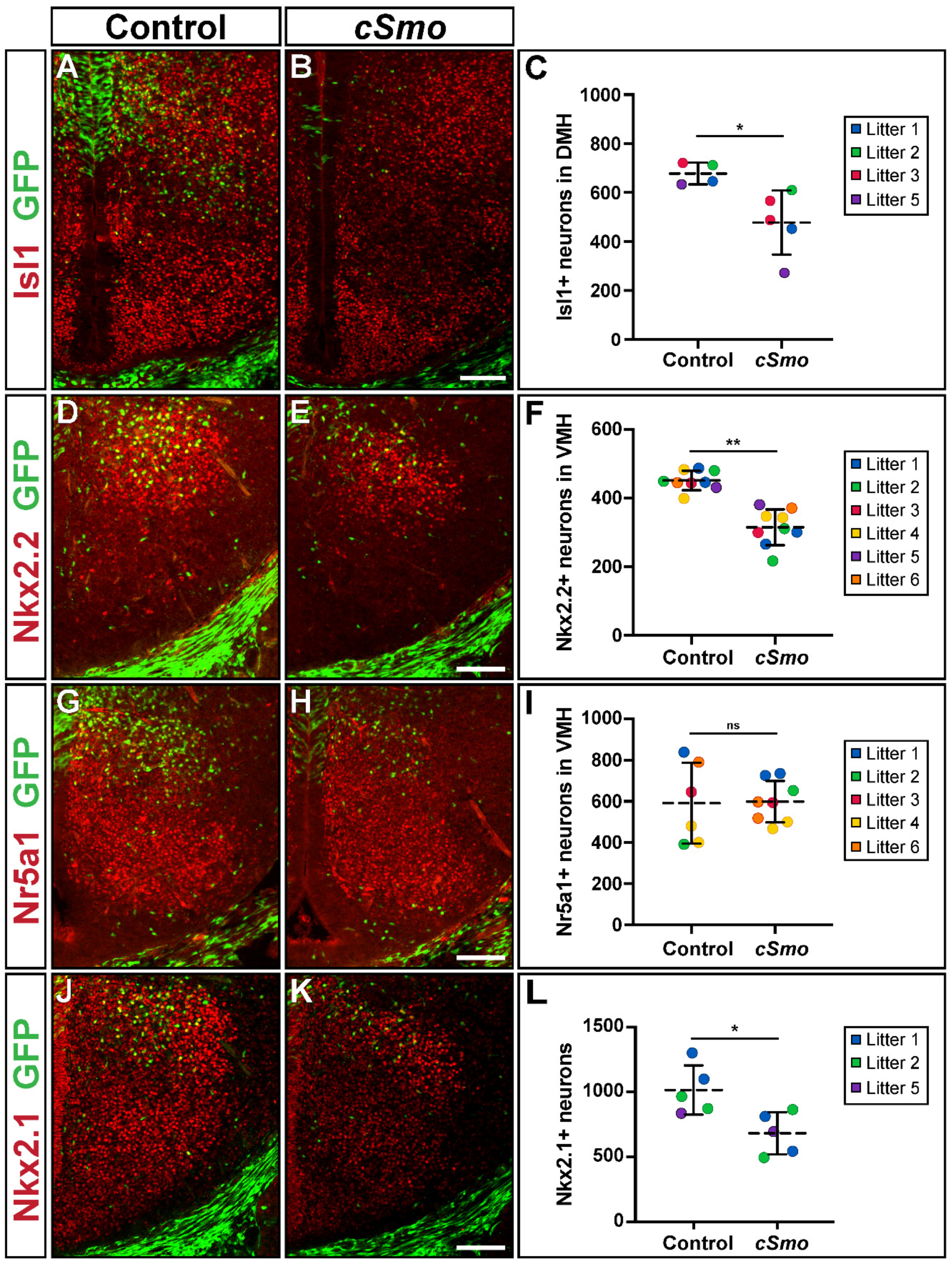
Shh signaling is required for subtype identity of VMH and DMH neurons. Coronal sections through the tuberal hypothalamus of control and *cSmo* embryos at E14.5 that received tamoxifen at E10.5. (A-C) The number of cells expressing Isl1 in the DMH is reduced in *cSmo* embryos (control n=4, *cSmo* n= 5, ^*^p=0.0229). (D-F) The number of post mitotic neurons expressing Nkx2.2 in the dorsal VMH is reduced in *cSmo* embryos compared to control littermates (n=9, ^**^p<0.0001). (G-I) The number of VMH neurons expressing Nr5a1 is equivalent between control and *cSmo* embryos (control n=6, *cSmo* n= 8, ns p>0.05). (J-L) The number of post-mitotic neurons expressing Nkx2.1 in the tuberal hypothalamus is reduced in *cSmo* embryos (n=5, ^*^p=0.0174). For all graphs, horizontal dotted line represents the mean and error bars indicate S.D. Each data point represents a single embryo that is color-coded by litter. Statistical analysis was performed using a two-tailed unpaired *t*-test.

### Non-cell autonomous defects in *cSmo* embryos alter VMH neuron subtype identity

Remarkably, despite the reduction of Nkx2.2 staining in *cSmo* embryos, the total number of Nr5a1 expressing neurons in the VMH was unchanged (Fig. 5G-I; *cSmo*: 598.5 ± 100.7, n=8, versus control: 590.8 ± 196.4, n=6, p=0.9253). Moreover, fewer Nr5a1 expressing neurons co-labeled with Nkx2.1 in *cSmo* embryos, which was especially notable in ventral regions of the VMH where *Gli1* is not normally expressed (Fig. 5J-L). The presence of defects in regions of the tuberal hypothalamus that are not known to be Shh responsive suggests that some of the phenotypes in *cSmo* embryos may occur through non-cell autonomous mechanisms.

The seemingly incongruous findings that Nkx2.2 neurons are partially reduced in the dorsal VMH of *cSmo* embryos without an effect on the total number of VMH cells prompted us to reassess the specification of tuberal hypothalamic progenitors. Nkx2.1 marks a broad ventral region of the ventricular zone in control embryos at E14.5, from the ventral midline of the hypothalamus to a dorsal limit that approximates the border between the VMH and DMH (Fig. 6A). Expression of Nkx2.2 overlaps with Nkx2.1 at the level of the dorsal VMH and extends dorsally into the ventricular zone of the prethalamus (Fig. 6A). There was a significant reduction in the number of cells expressing Nkx2.1 in the ventricular zone of *cSmo* embryos (Fig. 6A; *cSmo*: 131.0 ± 80.4, n=5, versus control: 259.4 ± 58.9, n=5, p=0.0205) and a concomitant ventral expansion of Nkx2.2 (Fig. 6A; *cSmo*: 118.2 ± 42.9, n=9, versus control: 58.0 ± 23.2, n=9, p=0.002). Given that Nkx2.2 is a direct transcriptional target of Shh signaling, it was puzzling that its expression was increased in *cSmo* mutants (Lei et al., 2006; Vokes et al., 2007). We evaluated other molecular readouts of Shh signaling and observed a similar ventral expansion of Olig2 (*cSmo*: 125.8 ± 12.9, n=5, versus control: 73.4 ± 6.5, n=5, p<0.0001) and Ki67 (*cSmo*: 107.0 ± 13.5, n=8, versus control: 81.8 ± 16.4, n=9, p=0.0037) in *cSmo* embryos (Fig. 6B,C). Notably, the cells displaying ectopic expression of Nkx2.2, Olig2 and Ki67 were not GFP positive, suggesting that they were wild type cells that had not undergone *Smo* recombination (Fig. S4). No change in the expression of *Gli1* was observed between *cSmo* and control embryos at E14.5, suggesting that the ectopic response to Shh signaling was transient and/or occurred at an earlier stage (Fig. 6D). The most parsimonious explanation for these results is that the conditional deletion of *Smo* prevented the normal uptake of Shh ligand, causing a non-cell autonomous gain in Shh signaling in neighboring wild type cells (Fig. 6E).

**Figure 6.**
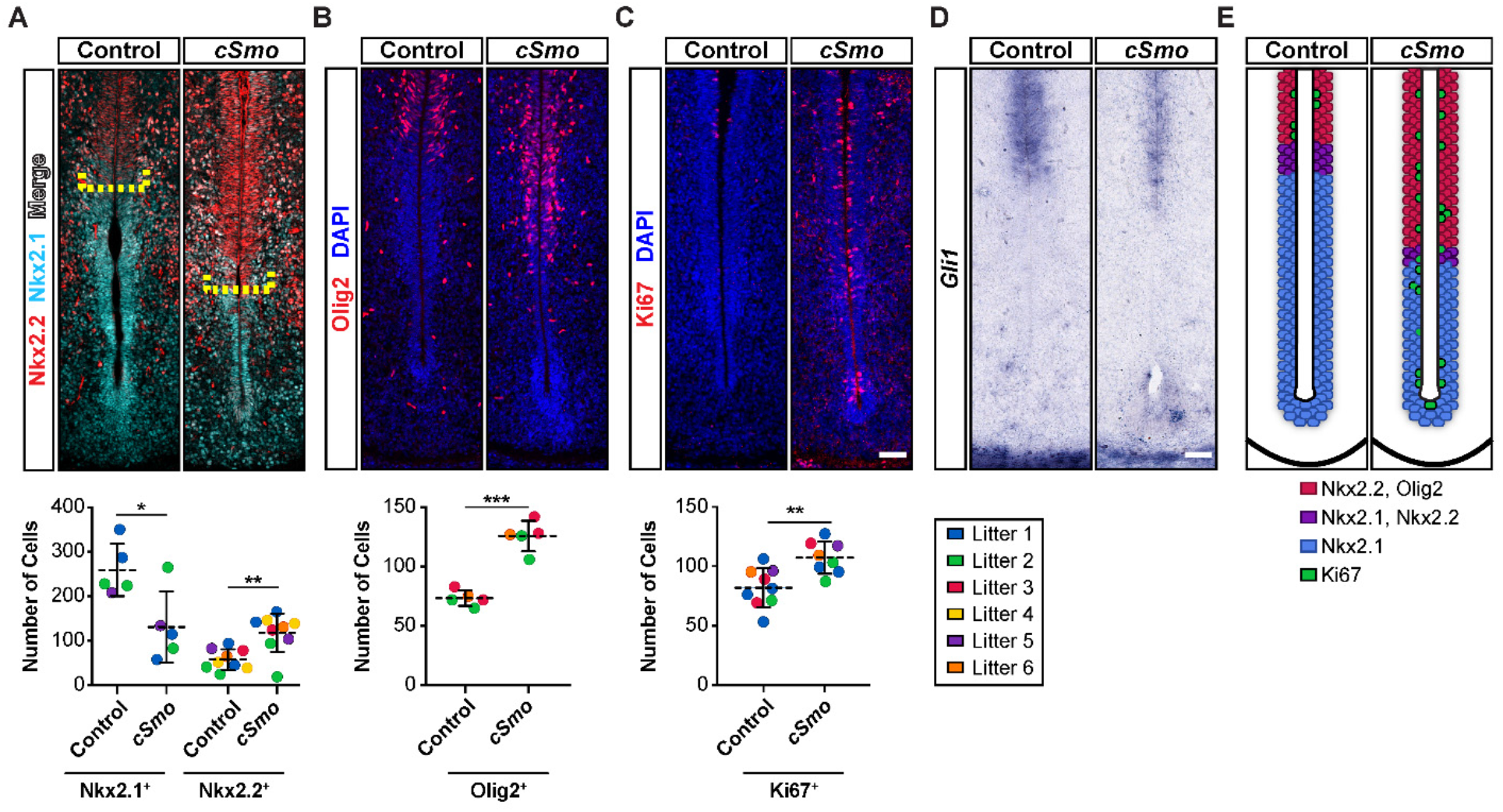
Non-cell autonomous alterations in hypothalamic progenitors in *cSmo* embryos. Coronal sections through the tuberal hypothalamus of control and *cSmo* embryos at E14.5 (tamoxifen administered at E10.5). (A) The boundary between Nkx2.1 and Nk2.2 expressing progenitors (yellow bracket) is shifted ventrally in the ventricular zone of *cSmo* embryos compared to control littermates. Ventral expansion in the number of Nkx2.2 expressing progenitors (n=9, ^**^p=0.002) is concomitant with a reduction in the number of cells expressing Nkx2.1 (n=5, ^*^p=0.0205). (B) The number of Olig2 expressing progenitors is ventrally expanded in *cSmo* embryos compared to control littermates (n=5, ^***^p<0.0001). (C) The number of Ki67 expressing progenitors is ventrally expanded in *cSmo* embryos compared to control littermates (control n=9, *cSmo* n=8, ^*\*\**^ p=0.0037). (D) *Gli1* expression is unchanged between control and *cSmo* embryos (n=3). (E) Schematic model depicting the non-cell autonomous gain in Shh responsiveness by ventral tuberal hypothalamic progenitors in *cSmo* mutant embryos compared to control littermates. For all graphs, horizontal dotted line represents the mean and error bars indicate S.D. Each data point represents a single embryo that is color-coded by litter. Statistical analysis was performed using a two-tailed unpaired *t*-test. Scale bars: 50 μm.

## DISCUSSION

Previous studies identified requirements for Shh at distinct spatial and temporal points in hypothalamic development, from induction to nucleogenesis (Chiang et al., 1996; Szabó et al., 2009; Shimogori et al., 2010; Zhao et al., 2012). Here, we further define the phenotypic consequences and pathogenic mechanisms attributed to loss of Shh signaling at early (E9.0) and late (after E10.5) stages of hypothalamic development. We determined that loss of hypothalamic nuclei in cases where Shh was deleted at early time points results from requirements for Shh in establishing proper dorsoventral patterning, neurogenesis, and limiting the expansion of the non-proliferative ventral midline. Using an inducible approach, we then characterized requirements for later Shh activity, at time points after dorsoventral patterning has been established, and found that Shh is necessary to maintain proliferation and subtype identity of VMH and DMH progenitors during peak periods of hypothalamic neurogenesis.

### Shh directs dorsoventral patterning and neurogenesis in the hypothalamus

Our results confirm that the early source of Shh in the hypothalamus is required to generate the full complement of tuberal hypothalamic neurons (Szabó et al., 2009; Shimogori et al., 2010; Zhao et al., 2012). In order to understand the mechanism underlying the loss of these cell types in *Shh^Δhyp^* mutants, we evaluated embryos at E10.5 and observed a failure to activate the appropriate patterning, proliferation and neurogenic programs. This result is consistent with the known morphogenetic role of Shh in establishing ventral neuronal identities through the regulation of homeodomain and bHLH transcription factors with some notable differences described below (Ericson et al., 1997; Briscoe et al., 1999; Briscoe et al., 2000; Muhr et al., 2001; Balaskas et al., 2012).

At spinal cord and hindbrain levels of the CNS, the notochord is the principal source of Shh required to specify ventral neuronal progenitors, whereas, the secondary source of Shh in the floor plate is needed for the formation of later born glial and ependymal cells types (Matise et al., 1998; Yu et al., 2013). In contrast to the notochord, expression of Shh in the prechordal plate is transient and only present for sufficient time to specify the hypothalamic territory, marked by Nkx2.1, but not the identity of neuronal progenitors (Szabó et al., 2009). Instead, it is the hypothalamic source of Shh that establishes VMH and DMH progenitor domains. Thus, whereas notochord derived Shh activates target genes, such as Nkx2.2 and Olig2, in the overlying neural tube, a similar function is fulfilled by the hypothalamic source of Shh. Given that hypothalamic neurons develop over a protracted period compared to most spinal cord neurons, it makes sense that they rely on an enduring source of Shh within the hypothalamus for their development (Davis et al., 2004; Altman and Bayer, 2006; Ishii and Bouret, 2012).

### Shh restricts ventral midline development in the tuberal hypothalamus

In addition to the dorsoventral patterning and neurogenic defects, our analysis of *Shh^Δhyp^* mutants revealed an unexpected expansion of ventral midline identity. It is likely that the increased number of ventral midline cells in *Shh^Δhyp^* embryos is a consequence of heightened Bmp signaling as evidenced by the expanded zone of non-proliferation and *Tbx3* expression, two molecular readouts of this pathway (Manning et al., 2006). Indeed, loss of Shh signaling in the anterior hypothalamus of *Shh^Δhyp^* embryos was previously shown to cause a rostral shift in *Bmp4* expression as early as E9.5 (Zhao, et al., 2012). Interestingly, a similar phenotype was also observed in *Lrp2^-/-^* mutant embryos, including downregulation of Shh signaling, flattened and expanded ventral midline, and a rostral shift in *Bmp4* expression (Christ et al., 2012). Lrp2 belongs to the LDL receptor gene family and binds both Shh (as a co-receptor) and Bmp4 (as a scavenger receptor) (McCarthy et al., 2002; Spoelgen et al., 2005; Christ et al., 2012). Therefore, a disruption in the balance of Shh and Bmp signaling, loss and gain respectively, is likely responsible for the ventral midline defects in *Lrp2^-/-^* and *Shh^Δhyp^* mutants.

A role for Shh in restricting ventral midline development in the tuberal hypothalamus is intriguing given the opposite function it plays in promoting floor plate induction in posterior regions of the CNS (Echelard et al., 1993; Roelink et al., 1995; Martí et al., 1995; Ericson et al., 1996; Chiang et al., 1996). Why might Shh have contrasting roles in the development of ventral midline cells at different levels of the neural tube? Firstly, the cells at the ventral midline of the tuberal hypothalamus, known as the median eminence, differ from floor plate cells in the spinal cord in that they are comprised of tanycytes, a specialized form of radial glia with neurogenic and gliogenic properties (Rizzoti and Lovell-Badge, 2017). In mammals, floor plate cells of the spinal cord are considered glial-like but do not give rise to neurons or glia. Tanycytes are an integral component of the hypophyseal portal system that connects the hypothalamus to the pituitary gland and regulate a variety of homeostatic processes given their access to blood borne signals (Rodríguez et al., 2005; Robins et al., 2013; Bolborea and Dale, 2013). Secondly, the hypothalamic ventral midline is a site of integration for multiple signaling pathways, including Shh, Bmp, Wnt and Fgf that help facilitate the unique organization of the hypothalamic-pituitary axis (Manning et al., 2006; Davis and Camper., 2007; Zhu et al., 2007; Potok et al., 2008; Zhao et al., 2012; Carreno et al., 2017; Fu et al., 2017). At the level of the spinal cord, Shh is the primary regulator of ventral neuronal identities, with additional signals serving to modulate Shh responsiveness (Gouti et al., 2015; Kong et al., 2015). Thirdly, Shh expressing cells in the hypothalamus are different from elsewhere in the neural tube in that they constitute actively dividing neuronal progenitors (Szabó et al., 2009; this study). Within the spinal cord, Shh expressing floor plate cells lack neurogenic properties. Given the clear differences in ventral development between the hypothalamus and spinal cord, it is not altogether surprising that Shh exhibits unique and overlapping functions at each of these CNS regions.

### Later Shh signaling- delineating fates

We sought to investigate roles and requirements for the enduring source of Shh in the tuberal hypothalamus. Our lineage tracing experiments revealed biased contributions of Shh expressing and Shh responding lineages to the VMH and DMH. On the basis of this observation, we wondered whether integration of progenitors may serve as a potential means by which neuronal heterogeneity is achieved within these hypothalamic nuclei.

Shh expressing and Shh responsive populations mix to give rise to the large domain of Nr5a1 cells in the VMH, though their contributions remain somewhat spatially discrete. Initially found broadly in the VMH, Nr5a1 expression becomes restricted to the VMH_DM_ by P0 (Cheung et al., 2013; Correa et al., 2015). Transcriptome analysis of the VMH in both neonatal and adult mice has revealed enrichment of 200 genes relative to expression in neighboring nuclei (Segal et al., 2005; Kurrasch et al., 2007). Examining the patterns of these markers revealed distinct spatial domains and biases, some overlapping with Nr5a1 and some mutually exclusive. Thus, it is clear that VMH neurons are diverse in nature. Molecular characterization of VMH neuronal subclasses has largely been performed in postnatal animals. Specifically, Islet1 and ERα are restricted to the VMH_VL_ (Davis et al., 2004), BDNF to the VMH_DM_ and VMH_C_ (McClellan et al., 2006), and leptin receptor to the VMH_DM_ (Balthasar et al., 2004; Dhillon et al., 2006). However, little is known regarding the processes through which these diverse populations are specified.Thus, within the VMH, the ventral bias of cells derived from Shh expressing progenitors and the dorsal bias of Gli1 expressing descendants may be indicative of a role for Shh signaling as a means of delineating distinct fates within a single nucleus.

The migration of a small number of Shh responsive progenitors into the VMH bears similarity to other hypothalamic cell types that originate from outside of their local progenitor territory. Most neuronal populations in the ARC are generated from multipotent progenitors that migrate radially from the adjacent ventricular zone (Li et al., 1996; McNay et al., 2006; Yee et al., 2009; Pelling et al., 2011; Lu et al., 2013; Lee et al., 2016), however some cells migrate tangentially from distal locations. For instance, GnRH neurons originate in the olfactory bulb and cross substantial distance to arrive at their final location within the ARC (Wray et al., 1989; Wray, 2002). Moreover, gene expression studies suggest that some Sst neurons migrate to the ARC from an anterior hypothalamic birthplace (Morales-Delgado et al., 2011). We propose that the migration of a Shh responsive cell population into the VMH, albeit from a nearby location, may further explain how cellular diversity is achieved in this nucleus.

### Later Shh signaling- maintaining proliferation of progenitor pools

In assessing the requirements of Shh at later stages of hypothalamic development, we observed a dependency on Shh for maintaining tuberal hypothalamic progenitors in a mitotically active state. When Shh signaling was disrupted after E10.5, there were fewer *cSmo* mutant cells in the VMH and DMH due to reduced cell division. Thus, Shh signaling is required to maintain hypothalamic progenitors in a mitotically active state during peak periods of VMH and DMH neurogenesis. These results are consistent with previous findings demonstrating a mitogenic role for Shh in other neurodevelopmental contexts (Rowitch et al.,1999; Wallace, 1999; Wechsler-Reya and Scott, 1999; Fuccillo et al., 2006; Komada et al., 2008).

### Non-cell autonomous phenotypes in *cSmo* mutants due to heightened Shh signaling

Our analysis of *cSmo* embryos also revealed that Shh is required to maintain the subtype identity of postmitotic VMH neurons. Despite the reduced number of Nkx2.2^+^, GFP^+^ neurons in the dorsal VMH of *cSmo* embryos, the total number of VMH neurons was unchanged. Moreover, fewer VMH neurons expressed Nkx2.1, especially in ventral regions where Smo was not deleted. We propose a non-cell autonomous mechanism to account for some of these differences in VMH neuron identity, whereby residual Shh ligand that was not taken up by *cSmo* mutant cells signaled to adjacent wild type progenitors, altering their patterns of gene expression and proliferation. In further support of this claim, a similar non-cell autonomous upregulation of Shh signaling was previously described upon the mosaic deletion of *Smo* in the ventral telencephalon (Xu et al., 2010).

Our observation of increased Shh signaling in non-recombined cells from *cSmo* mutant embryos is provocative in light of the association of somatic mutations in regulators of Shh signaling with hypothalamic hamartomas (Craig et al., 2008; Wallace et al., 2008; Hildebrand et al., 2016; Saitsu et al., 2016). Hypothalamic hamartomas are benign tumors that form during fetal brain development and, depending on their size and location, may disrupt endocrine function and cause seizures. Dominant mutations in *GLI3* that produce a truncated protein with constitutive repressor activity were identified in hypothalamic hamartomas (Craig et al., 2008; Wallace et al., 2008; Hildebrand et al., 2016; Saitsu et al., 2016). Individuals with Pallister Hall syndrome possess similar germline truncating mutations in *GLI3* and also develop hypothalamic hamartomas (Kang et al., 1997). Mutations in other components of the SHH signaling pathway, *OFD1* and *PRKRAC* (encoding the catalytic subunit ? of PKA), have also been described in resected hamartoma tissue (Hildebrand et al., 2016; Saitsu et al., 2016).

The mechanism of hypothalamic hamartoma formation is unknown. Paradoxically, the mutations that have been described in SHH pathway components are generally thought to inhibit SHH signaling, which, given its mitogenic activity, would be expected to suppress rather than promote the growth of this ectopic mass. Interestingly, the allele rate for somatic mutations in *GLI3* and *OFD1* ranged from 7-54%, suggesting that in some instances only a fraction of the cells in a given hypothalamic hamartoma may actually contain mutations (Saitsu et al., 2016). As with the *cSmo* mutants described in our study, somatic mosaicism in hypothalamic hamartomas may provide wild type cells with a temporary boost in SHH signaling and transient growth advantage, due to reduced ligand uptake by mutant cells that have lost their responsiveness to SHH. While experiments in our study did not extend into postnatal life and may not have targeted enough cells to induce hypothalamic hamartomas, they do nevertheless offer a compelling explanation for how these ectopic growths might form during brain development.

## MATERIALS AND METHODS

### Mice

All animal work was approved by the Institutional Animal Care and Use Committee (IACUC) at the Perelman School of Medicine, University of Pennsylvania. The *SBE2-cre* mouse line was previously described (Zhao, et al., 2012). *Gli1^CreER^* (*Gli1^tm3(cre/ERT2)Alj^*), *Gli1^lacZ^* (*Gli1 ^tm2Alj^*), *Rosa^ZsGreen^* (*Gt(ROSA)26Sor^tm6(CAG-ZsGreen1)Hze^*)*, Shh^+/−^*, *Shh^loxp/loxp^* (*Shh^tm2Amc^*), *Shh^CreER^* (Shh ^*tm2(cre/ERT2)Cjt*^), *Smo^+/-^* (*Smo^tm1Amc^*), and *Smo^loxp/loxp^* (*Smo^tm2Amc^*) mouse strains were procured from the Jackson labs (Bar Harbor, ME). Tamoxifen dissolved in corn oil was orally gavaged at 0.15 mg/g body weight to pregnant dams at appropriate developmental time points.

### Tissue dissection

Embryos were collected at the stated developmental age, with noon of plug day designated E0.5. Heads were dissected in cold PBS and fixed in 4% paraformaldehyde (PFA) for 90 minutes to overnight at 4°C. Heads were then cryoprotected in 30% sucrose and embedded in Tissue-Tek OCT Compound. Frozen tissue was cryosectioned at 16 μm (for E12.5 and E13.5 embryos), 18 μm (for E14.5 embryos), or 20 μm (for E10.5 and E18.5 embryos).

### Immunohistochemistry and section in situ hybridization

Immunohistochemistry was performed with the following antibodies: mouse anti-Ascl1 (1:100 BD Pharmingen, 556604), mouse anti-Gad1 (1:500, EMD Millipore, MAB5406), mouse anti-Isl1 (1:100, Developmental Studies Hybridoma Bank, 39.4D5), rabbit anti-Ki67 (1:1000, Novocastra. NCL-Ki67p), rabbit anti-Nkx2.1 (1:1000, Abcam, AB76013), mouse anti-Nkx2.2 (1:1000, DSHB, 74.5A5), rabbit anti-Olig2 (1:500, Millipore, AB9610), mouse anti-Pax6 (1:1000, Developmental Studies Hybridoma Bank, Pax6), rabbit anti-POMC (1:500, Phoenix Pharmaceuticals Inc., H-029-30), mouse anti-Nr5a1 (1:200, R&D Systems, PP-N1665), rat anti-Somatostatin (1:200, Millipore, MAB354) rabbit anti-Tyrosine Hydroxlase (1:1000, Pel-Freez, P40101-0). Somatostatin, Pax6, Nkx2.2, Nr5a1, Gad1, and Isl1 antibodies required antigen retrieval in 10mM citric acid buffer pH 6.0 at 90° C. Mouse on Mouse detection kit (Vector Laboratories BMK-2202) was used for blocking and primary antibody dilution of Nkx2.2, Gad1, and Isl1 antibodies.

Detection of primary antibodies was achieved using secondary antibodies conjugated to Cy3 anti-rabbit (1:200, 111-165-003, Jackson ImmunoResearch Laboratories), Cy3 anti-mouse (1:200, 115-166-006, Jackson ImmunoResearch Laboratories), Cy3 anti-rat (1:200, 112-166-003, Jackson ImmunoResearch Laboratories) or AlexaFluor 633 anti-rabbit (1:100, Invitrogen, A21070).

Section *in situ* hybridization with digoxygenin-UTP-labeled riboprobes was performed as described (Nissim et al., 2007). A minimum of three each of control and mutant embryos were evaluated for each antibody or *in situ* probe.

### β-galactosidase staining

Heads from *Gli1^lacZ/+^* embryos were dissected in cold PBS and fixed in 4% paraformaldehyde (PFA) for 90 minutes at 4°C. Heads were then cryoprotected in 30% sucrose and embedded in Tissue-Tek OCT Compound. Frozen tissue was cryosectioned at 20 μm. Slides were then stained in a solution containing 1 mg/ml X-gal at 37°C overnight. Following staining, slides were post-fixed in 4% PFA and washed in PBS.

### Quantification and statistical analysis

All cell counts were performed using the cell counter function in ImageJ (NIH) on tissue sections from at least three control and mutant embryos. In cases where double labeling was examined for GFP and another cell specific marker, the tissue was imaged at a single Z-plane. Each channel (green for GFP, red for the marker) was first examined independently, assigning a positive count for a given marker to the DAPI stained nucleus most closely associated with the staining. A cell was only counted as double labeled if a single nucleus marked by DAPI had been assigned to the cell labeled by GFP and the cell specific marker. Statistical analysis of all cell counts was performed in GraphPad Prism using the Student’s *t*-test.

## Acknowledgements

We thank Ishmail Abdus-Saboor, Sarah Millar, and Alex Rohacek for critical comments on the manuscript.

## Competing Interests

No competing interests declared.

## Author contributions

T.S.C. and S.E.Z. performed the experiments. T.S.C., S.E.Z and D.J.E. conceived the project.

T.S.C. and D.J.E. wrote the manuscript.

## Funding

This work was supported by a grant from the National Institutes of Health [R01 NS039421] to D.J.E

